# Amphitrophic *Listeria monocytogenes*: multi-dimensional genomic profiling reveals a third ecological strategy that challenges the virulence-persistence trade-off paradigm

**DOI:** 10.64898/2026.03.23.713700

**Authors:** Javier Gamboa

## Abstract

**Background:** The virulence-persistence trade-off is considered a fundamental organizing principle of *Listeria monocytogenes* population biology: hypervirulent clonal complexes dominate clinical cases but are rarely found in processing environments, while hypovirulent lineages dominate industrial niches but are underrepresented in severe disease. However, whether this dichotomy operates as an absolute paradigm has not been quantitatively evaluated at the population scale. Here we develop a multi-dimensional genomic scoring approach that simultaneously quantifies virulence potential (V), environmental persistence capacity (P), clonal epidemiological context (C), and antimicrobial resistance (R) across 903 genomes from four independent datasets spanning five countries, and apply it to test the universality of the trade-off and to characterize the ecological strategies of *L. monocytogenes* at the population level.

**Methods:** The scoring approach integrates four components into a composite 0–100 score through empirically calibrated weights (V: 30%, P: 40%, C: 20%, R: 10%). Validation employed 903 *L. monocytogenes* genomes from four public BioProjects: longitudinal industrial surveillance in Norway (Fagerlund et al. 2022, n = 513, PRJNA689484), retail environments in the United States (Stasiewicz et al. 2015, n = 191, PRJNA245909), clinical-environmental context in China (Wang et al. 2021, n = 151, PRJNA759341), and meat processing in Poland (Kurpas et al. 2020, n = 48, PRJNA629756).

**Results:** The composite score achieved excellent discriminatory performance for identifying persistent clones (AUC = 0.933; 95% CI: 0.910–0.954) with perfect specificity (1.000; zero false positives). The inverse V-P correlation was statistically significant across all four datasets (Spearman ρ from −0.144 to −0.713; p < 0.01), providing the first cross-dataset quantitative confirmation of the trade-off. However, simultaneous evaluation of V-P profiles at the population scale revealed that the species does not conform to a binary dichotomy but rather exhibits three quantitatively distinguishable ecological strategies, for which we propose a functional trophic taxonomy: *nosotrophic* lineages (22.7%; V > 65, P < 35), specialized in the pathogenic niche; *saprotrophic* lineages (5.8%; V < 30, P > 45), with irreversible virulence attenuation and industrial specialization; and, as the central finding, *amphitrophic* lineages (39.1%; V ≥ 35, P ≥ 40), which simultaneously retain functional *inlA* and stress tolerance determinants (SSI-1) without detectable genomic sacrifice. The three strategies differed significantly (Kruskal-Wallis H = 138.7; p = 7.6 × 10⁻³¹). The correspondence between trophic strategy and CC was predominant but not absolute, demonstrating that this phenotypic classification captures intra-CC functional heterogeneity inaccessible through conventional typing. Furthermore, comparison between genome-based and surveillance-informed classifications revealed that 60 hypervirulent isolates (CC1/CC14), genetically classified as nosotrophic, persisted for up to 8 years in industrial facilities despite lacking any recognized persistence markers — indicating that their prolonged survival reflects environmental opportunity rather than intrinsic genomic adaptation.

**Conclusions:** Multi-dimensional genomic profiling reveals that the virulence-persistence trade-off, while statistically robust, does not operate as an absolute paradigm. The amphitrophic strategy — documented here for the first time as a quantitatively distinguishable category encompassing 39.1% of the analyzed population — challenges the prevailing dichotomous model and identifies a previously unrecognized combined ecological niche. The ability to discriminate between genome-encoded persistence capacity and environmentally facilitated persistence provides a biological framework for understanding the ecological determinants of *L. monocytogenes* population dynamics in anthropogenic environments.

## 1. Introduction

### 1.1 Population biology of *L. monocytogenes*: the virulence-persistence trade-off

*Listeria monocytogenes* is an opportunistic intracellular pathogen responsible for listeriosis, with 20–30% case fatality rates in vulnerable populations (Charlier et al., 2017). Whole genome sequencing (WGS) has revealed a heterogeneous population structure stratified across multiple hierarchical scales, with the ecological dichotomy between clinical and environmental lineages representing one of the most striking features of the species’ biology.

Hypervirulent clonal complexes (CC1, CC2, CC4) dominate severe clinical cases (>70% of invasive listeriosis in French surveillance; Maury et al., 2016) but are rarely detected in processing environments (<15% of industrial isolates; Fagerlund et al., 2022). Conversely, hypovirulent lineages (CC9, CC121) dominate processing facilities (>60% of persistent isolates) but represent <15% of severe clinical cases (Maury et al., 2019). This ecological partition — the virulence-persistence trade-off — has a documented molecular basis: the PrfA/SigB regulatory switch mediates functional incompatibility between host invasion and environmental survival programs (Toledo-Arana et al., 2009).

However, two fundamental questions remain unresolved. First, while the transcriptomic model establishes incompatibility of *simultaneous expression* of virulence and saprophytic programs, it does not preclude *simultaneous carriage* of both genetic modules with conditional expression depending on ecological niche. The possibility that substantial proportions of the *L. monocytogenes* population maintain intact virulence and persistence arsenals has not been quantitatively evaluated at the population scale. Second, the degree of functional heterogeneity within individual clonal complexes — well documented at the genomic level through high-resolution cgMLST (Moura et al., 2016) — has not been systematically quantified through multi-dimensional profiling that integrates virulence, persistence, clonality, and resistance determinants simultaneously.

The present work addresses both questions through the development and validation of a multi-dimensional scoring approach that quantifies four independent biological dimensions across 903 genomes from four independent datasets, enabling the first population-scale assessment of ecological strategy diversity in *L. monocytogenes*.

### 1.2 Multi-scale biological complexity and intra-CC heterogeneity

The biological complexity of *L. monocytogenes* fundamentally limits assessment approaches based on single metrics. The species exhibits a heterogeneous population structure stratified across multiple hierarchical scales.

At the Clonal Complex (CC) level, the ecological dichotomy is well established. However, this dichotomy masks substantial intra-CC genetic heterogeneity revealed by high-resolution cgMLST typing (1,748 loci, Institut Pasteur scheme). Allelic distances (AD) of 5 to >150 between isolates of the same CC (Moura et al., 2016) reflect evolutionary divergence and differential ecological adaptation that CC classification collapses into a single categorical label. This intra-CC heterogeneity has direct functional implications: isolates of the same CC may differ in the presence of key virulence determinants (functional status of *inlA*, pathogenicity islands LIPI-3/LIPI-4) and persistence markers (biocide resistance genes, stress survival islets), generating a continuum of functional profiles inaccessible through categorical classifications. To date, no quantitative approach has systematically evaluated this intra-CC functional heterogeneity at the population scale. The present work documents for the first time the magnitude of this variability and its implications for understanding *L. monocytogenes* ecology.

### 1.3 The persistence paradox: absence of standardized criteria

The scientific community lacks formal standardized criteria for classifying bacterial persistence in industrial environments. As Ferreira et al. (2014) noted: “formal (statistical) criteria for the identification of persistence remain undefined.” Methodological heterogeneity manifests in temporal thresholds ranging from three months (Soumet et al., 2005) to several years (Fagerlund et al., 2022), and clonal identity criteria varying between ≤7 alleles (ECDC criterion) and ≤20 alleles (Fagerlund et al., 2022). Palaiodimou et al. (2021) confirm that the lack of a standardized definition hinders determination of true persistence.

There is furthermore a fundamental conceptual issue systematically overlooked in the literature: current persistence criteria are based exclusively on recurrent detection of the same clone over time, implicitly attributing prolonged survival capacity to intrinsic strain properties. However, recurrent detection does not discriminate between genuine persistence by biological merit of the isolate (genetic determinants for biocide resistance, stress tolerance, biofilm formation) and persistence by environmental opportunity — that is, strains without special adaptations surviving due to favorable ecological conditions in specific niches. This confusion between intrinsic capacity and environmental circumstance has direct biological implications: in the first case, persistence reflects adaptive evolution; in the second, it reflects niche availability independent of strain genotype.

The scoring approach described here explicitly addresses this distinction by separating the persistence component (P-Score) into two levels: Level 1 (genetic markers of intrinsic persistence capacity) and Level 2 (temporal evidence from longitudinal surveillance). This architecture enables differentiation between isolates with high genetic persistence potential that have not been detected longitudinally, and isolates without recognized persistence markers that persist due to environmental factors. Both scenarios generate recurrent detections indistinguishable by conventional criteria but reflect fundamentally different biological phenomena.

### 1.4 Existing approaches and their limitations

Several molecular typing systems currently operate for *L. monocytogenes*. Public schemes (ECDC/EFSA 1,748-locus cgMLST; CDC/GenomeTrakr) provide high-resolution clonal typing for outbreak tracing but do not integrate quantitative multi-dimensional profiling of virulence, persistence, and resistance into a unified assessment.

It is important to emphasize that cgMLST schemes fulfill with outstanding precision the function for which they were designed: determining the clonal identity of an isolate and its relatedness to other isolates for epidemiological traceability. The approach described here does not seek to replace this capability — it reuses cgMLST output as input for the clonality component and complements it with a fundamentally different question: what is the ecological profile of an isolate across multiple biological dimensions simultaneously.

### 1.5 The multi-dimensional scoring approach: definition and scope

We describe a multi-dimensional scoring approach that integrates four biological dimensions with robust empirical support in *L. monocytogenes* population biology into a composite quantitative score (0–100): (1) Clinical virulence potential (V-Score): genetic determinants associated with disease severity (Maury et al., 2016; Moura et al., 2016); (2) Environmental persistence capacity (P-Score): genetic adaptations and longitudinal detection patterns associated with prolonged survival in processing environments (Fagerlund et al., 2022; Møretrø & Langsrud, 2017); (3) Clonal epidemiological context (C-Score): position within the population structure through MLST/cgMLST typing (Moura et al., 2016; Ragon et al., 2008); and (4) Antimicrobial resistance burden (R-Score): genetic determinants of clinical resistance (Moura et al., 2024).

The approach implements dual weight calibration optimized for two analytical contexts: environment-focused (V:30%, P:40%, C:20%, R:10%) and clinical-focused (V:40%, P:20%, C:30%, R:10%). This calibration flexibility allows the same genomic data to be analyzed with weights reflecting the relative importance of each dimension in different ecological scenarios. The inclusion of the R-Score, despite currently low AMR prevalence in *L. monocytogenes* (<2% in Europe; Moura et al., 2024), provides integrated surveillance capacity for the documented emergence of antimicrobial resistance in foodborne pathogens (WHO GLASS, 2025; EFSA & ECDC, 2024) without affecting scoring performance when these determinants are absent, as demonstrated by the validation results.

## 2. Materials and Methods

### 2.1 Datasets

Validation employed 903 *L. monocytogenes* genomes from four public BioProjects, selected to represent geographic, sectoral, and epidemiological diversity (Table 1).

**Table 1.** Validation datasets.

| Dataset | BioProject | N | Context | Country | Period | Reference |
| --- | --- | --- | --- | --- | --- | --- |
| Fagerlund | PRJNA689484 | 513 | Industrial, 15 processing plants | Norway | 1990–2020 | Fagerlund et al., 2022 |
| Stasiewicz | PRJNA245909 | 191 | Retail delis, 3 states | USA | 2010–2012 | Stasiewicz et al., 2015 |
| Wang | PRJNA759341 | 151 | Clinical + environmental | China | 2014–2018 | Wang et al., 2021 |
| Kurpas | PRJNA629756 | 48 | RTE meat + environment | Poland | 2014–2017 | Kurpas et al., 2020 |
| Total |  | 903 |  | 5 countries, 3 continents |  |  |

The Fagerlund dataset constitutes the primary development and validation dataset, providing unprecedented temporal and spatial documentation: 30 years of continuous surveillance across 15 processing facilities (9 meat, 6 salmon) with factory, isolation year, and clonal identity (SNP reference) metadata enabling reconstruction of persistence patterns at the individual clone level. The three remaining datasets provide cross-validation in independent geographic and epidemiological contexts.

### 2.2 Bioinformatic pipeline

All genomes were processed through a standardized pipeline. Quality control was performed with FastQC (coverage ≥30×, Q30 ≥80%), followed by *de novo* assembly with SPAdes v3.15. MLST typing (7 housekeeping loci: *abcZ*, *bglA*, *cat*, *dapE*, *dat*, *ldh*, *lhkA*) was performed against the Institut Pasteur database (PubMLST) for Sequence Type (ST) and Clonal Complex (CC) assignment. cgMLST typing employed chewBBACA v3.1.2 with the Institut Pasteur 1,748-locus scheme for allelic distance (AD) calculation and hierarchical clustering (HC). Virulence marker detection (LIPI-1/3/4, *inlA*) was performed using ABRicate with the VFDB database, biocide resistance gene identification (*qacH*, *bcrABC*, SSI-1) via BLAST against reference sequences, and AMR determinant detection through a tripartite pipeline (CARD v3.2.4, ResFinder 4.1, NCBI AMRFinderPlus v3.11). Detection thresholds were ≥90% identity and ≥80% coverage for all markers, except *qacEΔ1* (≥95% identity, ≥90% coverage).

### 2.3 Component V: Virulence Potential (0–100)

The V-Score assesses the potential to cause severe invasive listeriosis through hierarchical integration of three levels:

Level 1: Phylogenetic context (0–40 points). CC classification provides the primary population context. Isolates are stratified into three categories based on association with documented severe listeriosis in European surveillance (Maury et al., 2016; n = 6,633 isolates): hypervirulent (40 pts: CC1, CC2, CC4, CC14, CC87), intermediate (20 pts: CC3, CC5, CC6, CC7, CC11), and hypovirulent (5 pts: CC9, CC121, and remaining). CC6 classification as intermediate is based on recent evidence documenting average virulence for ST6. CC14 and CC87 are classified as hypervirulent based on their documented association with severe outbreaks. Genetic proximity to documented clinical lineages (minimum AD in cgMLST against NCBI Pathogen Detection and Institut Pasteur BIGSdb reference databases) adds up to 10 additional points, capturing intra-CC heterogeneity.

Level 2: Pathogenicity islands (−10 to +35 points). LIPI-1, universally conserved, is penalized (−10 pts) only if deletions are present in key genes (*prfA*, *hly*, *actA*). LIPI-3 (+15 pts) is scored exclusively in a Lineage I context, where its presence has validated functional significance (association with neurolisteriosis; Cotter et al., 2008). LIPI-4 (+20 pts) is restricted to CC4/CC87/CC382/CC619, based on restricted distribution and association with maternal-neonatal listeriosis (Maury et al., 2016; clinical frequency 71.3%, p < 10⁻⁴). Detection: ≥7/8 genes for LIPI-3 (≥90% id, ≥90% cov) and ≥5/6 genes for LIPI-4 (≥85% id, ≥80% cov).

Level 3: *inlA* functionality (−40 to +20 points). Functional status is determined through *in silico* extraction and translation of the coding sequence, PMSC search, and alignment against EGD-e reference (800 aa). Scoring: complete InlA (≥780 aa, no PMSCs): +20 pts; truncated (PMSC at intermediate position): −15 pts; severely truncated (<500 aa): −40 pts. The invasiveness reduction associated with PMSCs (10–100× in murine models; Nightingale et al., 2005) and the differential prevalence between lineages (40–50% in CC9/CC121 vs <5% in CC1/CC4; Van Stelten et al., 2010) underpin this weighting.

Normalization: V_raw is scaled to 0–100. Negative values (severely attenuated isolates) are adjusted to zero.

### 2.4 Component P: Persistence Risk (0–100)

The P-Score integrates genetic markers of environmental adaptation (Level 1) with temporal and spatial detection patterns (Level 2), explicitly distinguishing between intrinsic persistence potential and longitudinal evidence of observed persistence.

Level 1: Genetic markers (0–75 points). Three categories of determinants are evaluated:

Disinfectant resistance: *qacEΔ1* (+15 pts; truncated QAC efflux pump, high prevalence in persistent strains), *qacH* (+10 pts; complete QAC efflux pump encoded on Tn6188), complete *bcrABC* operon (+20 pts; ABC transporter system for benzalkonium chloride resistance; all three *bcrA/B/C* genes must be detected at ≥90% id, ≥85% cov).

Stress tolerance: SSI-1 (+15 pts; ~9 kb stress survival islet including glutamate decarboxylase system *gadD/E/T* and *sigB* factor; detection of ≥6/9 genes required), cadmium tolerance (+5 pts; *cadA/cadC* cassette detection), absence of CRISPR-Cas system (+5 pts; facilitates adaptive genetic material uptake).

Genomic adaptation factors: Complete flagellar motility, cold shock genes, oxidative tolerance, prophage presence — contribute to overall adaptation score.

Level 2: Temporal-spatial patterns (0–50 points). When longitudinal surveillance metadata are available, three metrics are quantified:

- Colonization time (0–30 pts): Difference between last and first detection of the same clone (≤7 AD cgMLST) in the same facility. <4 weeks: 0 pts; 4–12 weeks: 10 pts; 13–52 weeks: 20 pts; >52 weeks: 30 pts.
- Detection frequency (0–10 pts): Number of independent detections. <3: 0 pts; 3–4: 5 pts; ≥5: 10 pts.
- Spatial dispersion (0–10 pts): Number of facilities where the clone has been detected. 1: 0 pts; 2: 5 pts; ≥3: 10 pts (pervasive clone *sensu* Fagerlund et al., 2022).

When temporal metadata are unavailable, Bayesian imputation is applied: E[T|G] = (G/G_max) × 25, where G is the Level 1 genetic score. This conservative estimator positions the score in the intermediate range between absence of persistence and maximum confirmed persistence.

Normalization: P_raw / 75 × 100, capped at [0, 100].

### 2.5 Component C: Clonality Context (0–100)

The C-Score quantifies the epidemiological context of the isolate through two levels that evaluate, respectively, genetic proximity to known surveillance populations and the clonal expansion success of the lineage. Unlike the V-Score — which employs cgMLST proximity specifically against clinical isolates to infer virulence potential — the C-Score evaluates the isolate’s position in the overall epidemiological structure (clinical + environmental + outbreak).

Level 1: Clonal genetic proximity (0–70 points). The minimum allelic distance (AD_min) between the isolate and its nearest neighbor in reference surveillance databases (NCBI Pathogen Detection, n > 30,000 genomes; Institut Pasteur BIGSdb, n > 15,000) is calculated. Typing employs the 1,748-locus cgMLST scheme via chewBBACA v3.1.2 (quality criterion BSR ≥ 0.6). Scoring thresholds derive from international surveillance agency consensus:

- AD ≤ 4: 70 pts (ECDC multi-country threshold for cross-border outbreaks)
- AD ≤ 7: 60 pts (Institut Pasteur Cluster Type)
- AD ≤ 10: 45 pts (Ruppitsch et al., 2015 criterion)
- AD ≤ 20: 30 pts (extended outbreak range)
- AD ≤ 50: 18 pts; AD ≤ 100: 10 pts; AD ≤ 150: 5 pts (Sublineage boundary)
- AD > 150: 0 pts (no epidemiologically relevant clonal relationship)

Level 2: Clonal expansion characteristics (0–30 points). Two complementary metrics of lineage evolutionary success are evaluated:

Metric 2.1 — HC10 cluster size (0–20 pts): Member count of the cluster defined at AD ≤ 10 threshold in the surveillance database. Reflects clone prevalence and geographic distribution. ≥100 isolates: 20 pts (epidemic clone); 50–99: 15 pts; 20–49: 10 pts; 5–19: 5 pts; 2–4: 3 pts; singleton: 0 pts.

Metric 2.2 — Cluster genetic compactness (0–10 pts): AD_median / AD_max ratio within the HC10 cluster. Evaluates temporal expansion dynamics: genetically compact clusters (ratio > 0.5; +10 pts) indicate recent and active expansion, while dispersed clusters (ratio < 0.3; 0 pts) indicate accumulated evolutionary diversification during prolonged circulation. For clusters with < 5 isolates, AD_max is used as proxy (≤10: +10 pts; 11–30: +5 pts; >30: 0 pts).

Contingency mechanism: if the cgMLST profile is unavailable (insufficient quality, <95% locus coverage), CC representation in PubMLST is used as proxy.

### 2.6 Component R: Antimicrobial Resistance (0–100)

The R-Score quantifies the antimicrobial resistance burden of the isolate, evaluating both the presence of specific genetic determinants and the epidemiological relevance of their transferability. The two-level structure distinguishes chromosomal from transferable resistance, recognizing that detection of resistance to first-line therapeutic agents (ampicillin-gentamicin) confers the highest priority.

Level 1: Critical genetic determinants (0–80 points). Four categories are evaluated, stratified by clinical importance:

Metric 1.1 — β-lactam resistance (ampicillin): Detection of β-lactamases (*blaZ*, *blaT*, *blaL*); +5 pts/gene. Critical given that ampicillin is first-line treatment for invasive listeriosis (prevalence <5% in European surveillance; EUCAST 2023).

Metric 1.2 — Aminoglycoside resistance (gentamicin): Detection of bifunctional enzyme *aac(6’)-Ie-aph(2’’)-Ia*; +10 pts. Essential for its role in ampicillin-gentamicin combination therapy for CNS listeriosis and endocarditis.

Metric 1.3 — Tetracycline resistance: Detection of *tetM*, *tetS*, *tetL*, *tetK* genes; +10 pts/gene (maximum 40 pts). Indicator of emerging resistance (Moura et al., 2024).

Metric 1.4 — Fluoroquinolone resistance (QRDR mutations): Substitutions in *gyrA* (positions Ser98, Asp105) and *parC* (Ser83, Glu87), EGD-e reference; +15 pts/mutation (maximum 60 pts).

Detection: BLAST ≥90% identity, ≥80% coverage against CARD/AMRFinderPlus.

Level 2: Multi-resistance megaplasmids (0–20 points). +20 pts are assigned if a plasmid ≥40 kb carrying ≥3 AMR genes from different categories is detected, reflecting the biological significance of horizontal transfer of multiple resistance in a single conjugation event.

Regional amplification factor: F = max(1.0; P_ref / P_AMR) where P_ref = 5%. In regions with ultra-low AMR prevalence (<1%), resistance detection represents an exceptional event justifying score amplification. In regions with endemic AMR (>10%), the factor is neutral. R_final = min(100, R_raw × F_regional).

### 2.7 Weighted integration and contextual calibration

The composite score is calculated as a weighted sum:

Score = V × w_V + P × w_P + C × w_C + R × w_R

Weights depend on the analytical context:

Environment-focused calibration (validated in this work): w_V = 0.30; w_P = 0.40; w_C = 0.20; w_R = 0.10. Prioritizes persistence as the primary ecological driver, grounded in the principle that in processing environments, chronic colonization (persistence) dominates the biological dynamics over individual virulence events.

Clinical-focused calibration: w_V = 0.40; w_P = 0.20; w_C = 0.30; w_R = 0.10. Prioritizes virulence (clinical severity) and clonality (epidemiological traceability).

### 2.8 Validation strategy

Layer 1 — Primary development and validation (Fagerlund, n = 513). Clonal persistence is defined as recurrent detection of the same clone (≤7 AD cgMLST, shared SNP reference) in the same facility across ≥2 distinct calendar years. Temporal metadata (factory, year) were extracted from the supplementary material of Fagerlund et al. (2022) and integrated with composite scores to compute performance metrics.

Layer 2 — Geographic cross-validation (Stasiewicz + Wang + Kurpas, n = 390). Verification of V-P trade-off consistency, trophic strategy pattern reproducibility, and inter-component correlation stability.

Layer 3 — Independence and robustness analysis. Inter-component Spearman correlations, per-component AUC and Cohen’s d, and intra-CC heterogeneity quantification.

### 2.9 Statistical analysis

All analyses were performed in Python 3.10 with scipy v1.11.2, scikit-learn v1.3.0, numpy v1.24, and pandas v2.0. 95% CIs for AUC estimated via bootstrap (2,000 resamples). Optimal classification threshold by Youden index. Effect size by Cohen’s d.

### 2.10 Functional trophic taxonomy: definitions

Based on the quantitative V-P profiles observed at the population scale, this work proposes a tripartite functional taxonomy of *L. monocytogenes* ecological strategies across the host–industrial environment continuum. The proposed terms derive from Greek roots describing the primary trophic niche (*trophe* = nourishment, sustenance in a niche):

- *Nosotrophic* (*nosos* = disease + *trophe*): lineage whose fitness is primarily expressed in the pathogenic niche. High virulence, low environmental persistence.
- *Saprotrophic* (*sapros* = decomposition + *trophe*): lineage whose fitness is primarily expressed in the saprophytic niche. Virulence irreversibly attenuated (PMSCs in *inlA*), high environmental persistence.
- *Amphitrophic* (*amphi* = both + *trophe*): lineage whose fitness is expressed in both niches without detectable genomic sacrifice. Retains functional virulence (intact *inlA*) alongside environmental tolerance determinants (SSI-1). The term evokes the analogy with amphibious organisms, capable of alternating between habitats.

The quantitative assignment criteria are phenotypic (based on V and P scores calculated for each individual isolate, not on CC membership):

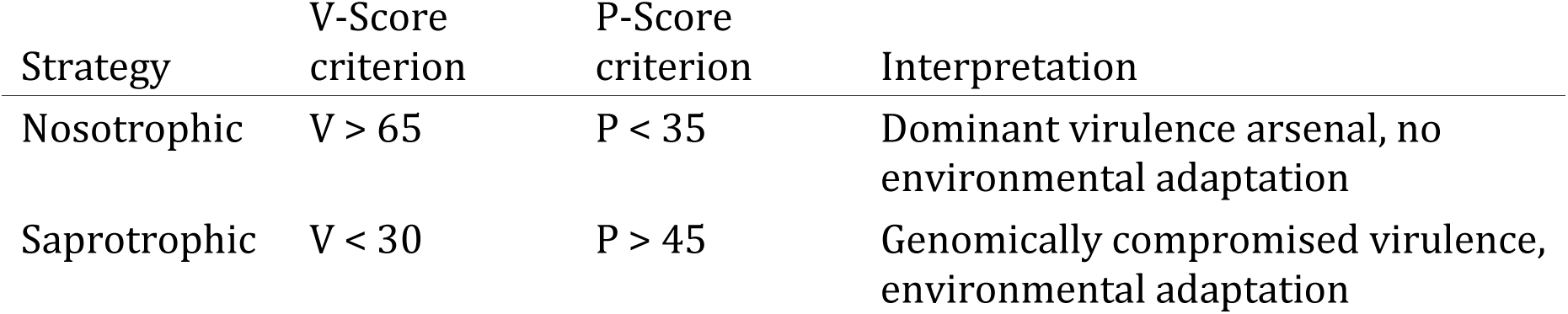
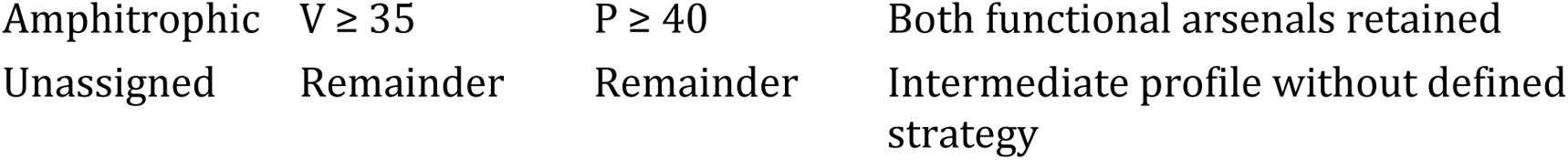

It is important to note that this classification operates on the quantitative functional profile, not on the CC categorical label. A CC121 isolate with complete *inlA* (V ~ 47) and SSI-1 (high P) would be classified as amphitrophic, not saprotrophic, regardless of the majority of its CC exhibiting the opposite pattern. This intra-CC resolution is precisely the discriminatory capability that the multi-dimensional approach contributes beyond CC classification.

## 3. Results

### 3.1 Discriminatory performance of the composite score

The composite score, calculated with environment-focused calibration (V × 0.30 + P × 0.40 + C × 0.20 + R × 0.10), demonstrated excellent discriminatory performance for identifying persistent clones in the Fagerlund dataset (Figure 1, Figure 4; Table 2).

**Table 2.**
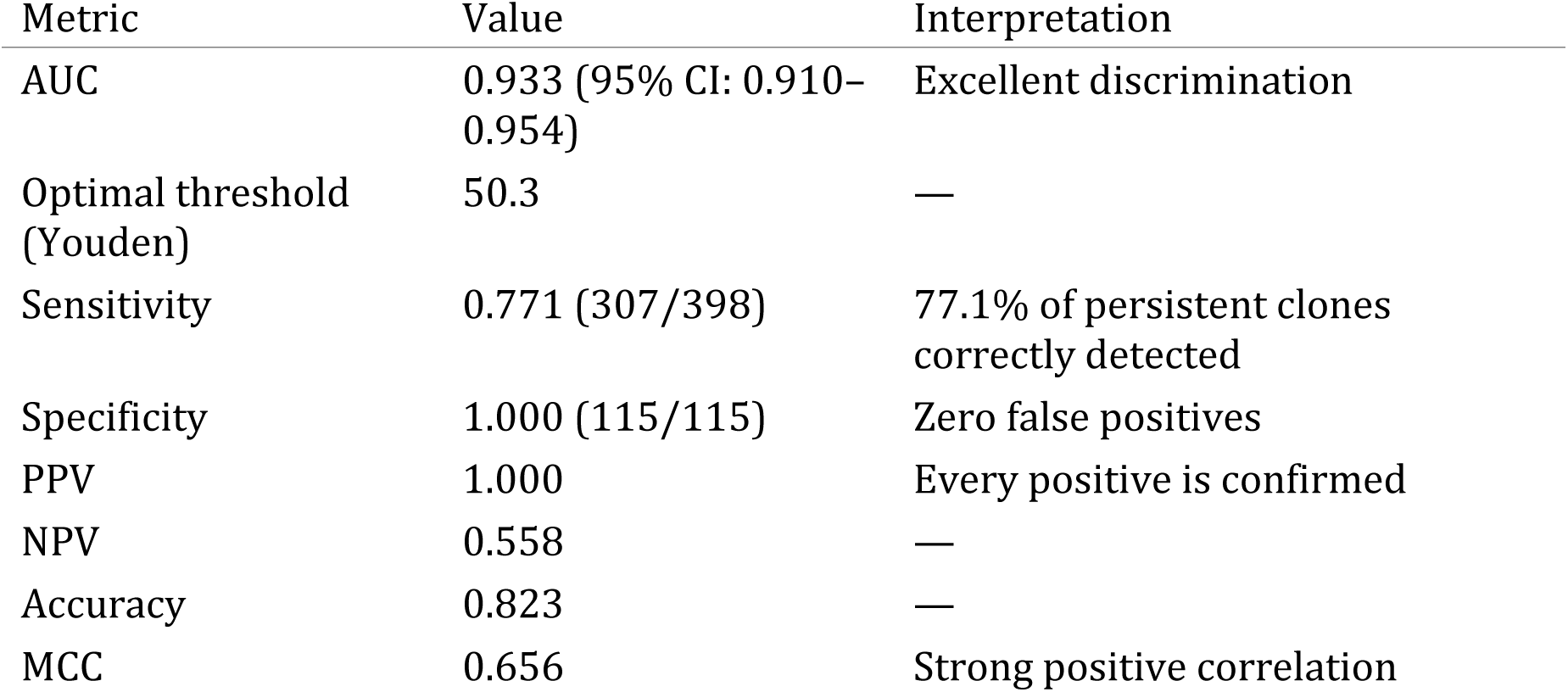
Performance metrics (Fagerlund, n = 513)

The perfect specificity (1.000) indicates that no sporadically detected isolate was incorrectly classified above the threshold. The sensitivity of 0.771 indicates that 22.9% of persistent clones received scores below the threshold, corresponding to clones with atypical genomic profiles whose persistence likely depends on environmental factors (facility configuration, niche availability) rather than intrinsic genomic adaptations — precisely the distinction that the P-Score’s bi-level architecture is designed to capture.

### 3.2 Differential component contribution

The P-Score achieved AUC = 0.980 (Figure S1) (Cohen’s d = +3.22, enormous effect), confirming persistence markers as the component with highest informative load in the environmental context. The V-Score exhibited AUC = 0.446 (<0.5), indicating that higher-virulence isolates tend to be sporadically detected — empirical validation of the virulence-persistence trade-off.

**Table 3.** Per-component performance (Fagerlund, n = 513)

| Component | AUC | Cohen's d | p-value | Confirmed biological role |
| --- | --- | --- | --- | --- |
| V-Score | 0.446 | -0.34 | 0.076 | Inverse correlation: validates trade-off |
| P-Score | 0.980 | +3.22 | $8.8 \times 10^{-56}$ | near-perfect discrimination |
| C-Score | 0.523 | +0.03 | 0.448 | Neutral: dataset clonal homogeneity |
| R-Score | 0.501 | +0.01 | 0.929 | Neutral: low AMR prevalence in dataset |

The C and R components showed AUC values near 0.5 (no discriminatory contribution), reflecting dataset-specific characteristics: clonal homogeneity (dominated by CC9/CC121/CC7) and low AMR prevalence (<1%). This behavior is consistent with the scoring design: C-Score and R-Score contribute discrimination when signal exists (clonal diversity, emerging AMR) and remain neutral — without generating noise — when that signal is absent. In contexts with greater clonal diversity (multi-sector surveillance) or emerging AMR prevalence, both components would contribute substantial discrimination.

### 3.3 Virulence-persistence trade-off quantified cross-dataset

The inverse V-P correlation was confirmed as statistically significant across all four datasets (Table 4, Figure S2), constituting the first cross-dataset empirical quantification of this phenomenon through integrated genomic scoring.

**Table 4.**
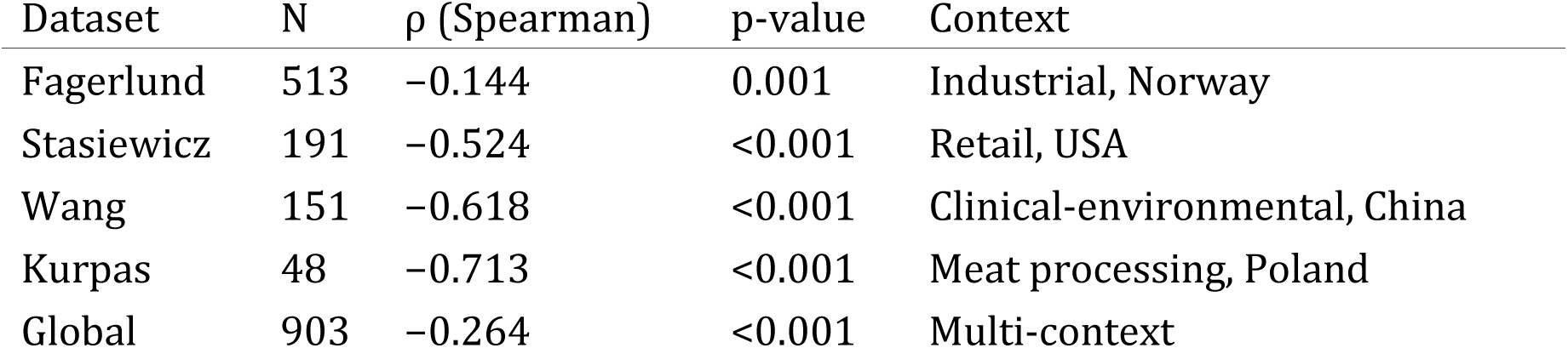
V-P trade-off across four independent datasets.

The direction of the effect was uniform, confirming that the virulence-persistence trade-off constitutes a robust biological phenomenon reproducible across heterogeneous geographic and epidemiological contexts.

### 3.4 Informative independence between components

Five of six pairs exhibited |ρ| < 0.25, confirming substantial independence (Figure 3). In the global analysis (N = 903), the P-R correlation reached ρ = +0.591 (p < 0.001), indicating co-selection between persistence and antimicrobial resistance — a finding with implications for understanding the ecology of resistance dissemination.

**Table 5.** Spearman correlation matrix (Fagerlund, n = 513)

|  | V | P | C | R |
| --- | --- | --- | --- | --- |
| V | 1.000 | -0.144*** | -0.235** | * -0.093* |
| P |  | 1.000 | -0.022 | +0.148*** |
| C |  |  | 1.000 | +0.024 |
| R |  |  |  | 1.000 |

### 3.5 Three trophic strategies: an emergent functional taxonomy

Simultaneous evaluation of V-Score and P-Score at the population scale revealed that the structure of *L. monocytogenes* does not conform to the virulent-vs-persistent dichotomy assumed as paradigm, but rather exhibits three quantitatively distinguishable ecological strategies (Table 6, Figure 2). We propose for these three strategies the functional taxonomy defined in section 2.10, based on the primary trophic niche of each group.

**Table 6.** Trophic strategies of *L. monocytogenes* identified by V-P phenotypic classification (N = 903)

| Strategy | Criterion | N | % | V | P | Score | InlA intact | SSI-1 | qacH |
| --- | --- | --- | --- | --- | --- | --- | --- | --- | --- |
| Nosotrophic | V>65,<br>P<35 | 205 | 22.7 | 75.4<br>± 6.4 | 22.9<br>± 1.1<br>4 | 2.6 1 | 00%<br>0 | % 0 | % |
| Amphitrophic | V≥35,<br>P≥40 | 353 | 39.1 | 56.5<br>± 6.8 | 55.8<br>±<br>13.6<br>4 | 5.1 1 | 00%<br>8 | 7.3%<br>0 | % |
| Saprotrophic | V<30,<br>P>45 | 52 | 5.8 | 20.3<br>± 6.3 | 63.6<br>±<br>12.4<br>3 | 3.0 0 | % 9 | 0.4%<br>4 | 0.4% |
| Unassigned | Remainder | 293 | 32.4 | 36.1<br>± | 31.3<br>± | 9.6 6 | 3.5%<br>2 | 5.6%<br>8 | .2% |

| Strategy | Criterion | N | % | V | P | Score | InlA<br>intact | SSI-1 | qacH |
| --- | --- | --- | --- | --- | --- | --- | --- | --- | --- |
|  |  |  |  | 19.1 | 13.6 |  |  |  |  |
|  |  |  |  |  | 2 |  |  |  |  |

Kruskal-Wallis: $H = 138.7$ ; $p = 7.6 \times 10^{-31}$ . All pairwise comparisons significant (Mann-Whitney, $p < 0.001$ ).

It is relevant to note that the genomic marker profiles observed in each category are a direct consequence of the scoring structure, not a coincidence: the V > 65 threshold requires functional *inlA* (contributing +20 pts) plus hypervirulent phylogenetic context (+40 pts), so every nosotrophic isolate necessarily carries complete *inlA*. Conversely, V < 30 is only achievable with penalties for truncated (−15) or severely truncated *inlA* (−40), confirming that every saprotrophic isolate exhibits irreversible loss of InlA functionality. Amphitrophic isolates (V ≥ 35, P ≥ 40) require both functional *inlA* and environmental tolerance markers, and indeed 100% retain complete *inlA* while 87.3% carry SSI-1. This coherence between quantitative thresholds and underlying biology validates the scoring structure.

The correspondence between trophic category and CC is not absolute but predominant — precisely the distinction the multi-dimensional approach contributes beyond categorical classifications (Table 7).

**Table 7.**
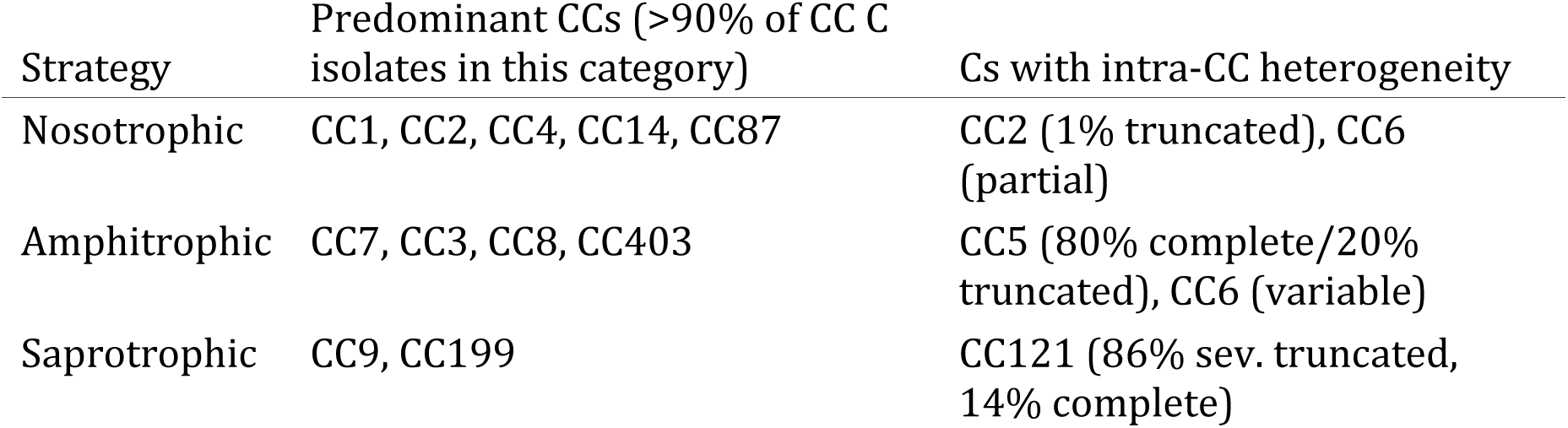
Correspondence between phenotypic trophic strategy and predominant CCs Predominant CCs (>90% of CC C.

| Strategy | Predominant CCs (>90% of CC C isolates in this category) | Cs with intra-CC heterogeneity |
| --- | --- | --- |
| Nosotrophic | CC1, CC2, CC4, CC14, CC87 | CC2 (1% truncated), CC6 (partial) |
| Amphitrophic | CC7, CC3, CC8, CC403 | CC5 (80% complete/20% truncated), CC6 (variable) |
| Saprotrophic | CC9, CC199 | CC121 (86% sev. truncated, 14% complete) |

The existence of CCs with isolates distributed across multiple trophic categories (Figure 5) — particularly CC5, CC6, and CC121 — demonstrates that CC classification is insufficient to capture actual functional diversity. A CC121 isolate with complete *inlA* (14% of isolates in this CC) is not functionally equivalent to a CC121 with severely truncated *inlA*, and the multi-dimensional approach classifies them into distinct trophic categories. This intra-CC resolution constitutes a discriminatory advantage over categorical approaches.

Nosotrophic lineages (V > 65, P < 35; n = 205): Specialized in the pathogenic niche. Functional *inlA* and hypervirulent phylogenetic context required by the V threshold. No SSI-1 or biocide resistance markers. Predominate in CC1, CC2, CC4, CC14, and CC87 — the classic hypervirulent lineages that dominate severe invasive listeriosis and exhibit transient presence in industrial facilities.

Saprotrophic lineages (V < 30, P > 45; n = 52): Specialized in the saprophytic niche. Genomically compromised virulence — the V < 30 threshold necessarily requires truncated or severely truncated *inlA*, confirming irreversible loss of InlA functionality. Carry *qacH* (40.4%) and SSI-1 (90.4%). Predominate in CC9 and CC199. They have performed the trade-off irreversibly: PMSCs in *inlA* constitute a permanent genomic loss of invasive capacity, consistent with the paradigm established by Maury et al. (2016, 2019).

Amphitrophic lineages (V ≥ 35, P ≥ 40; n = 353): The central finding of this work. They represent 39.1% of the analyzed population and constitute, to our knowledge, the first quantitatively distinguishable category at the population level that escapes the dichotomous paradigm of the virulence-persistence trade-off. The thresholds simultaneously require functional *inlA* (V ≥ 35) and environmental tolerance markers (P ≥ 40), with verified moderate-to-high virulence (V = 56.5; 100% functional InlA) and elevated persistence (P = 55.8; 87.3% SSI-1). Their persistence strategy relies on stress tolerance (SSI-1: acid, salt, and oxidative tolerance) without the specific quaternary disinfectant resistance (*qacH*, *bcrABC*) that characterizes saprotrophic lineages. They predominate in CC7, CC5, CC3, CC8, CC403, and CC6, but assignment is individual — atypical isolates within these CCs would fall into other categories.

The amphitrophic pattern was consistent across all four datasets: 29.4% in Fagerlund, 69.6% in Stasiewicz, 41.7% in Wang, and 12.5% in Kurpas, confirming its reproducibility across heterogeneous epidemiological contexts.

### 3.6 Documented intra-CC heterogeneity

The multi-dimensional approach demonstrated the ability to differentiate isolates within the same CC. CC121 (n = 86): 80/86 isolates exhibited severely truncated *inlA* (V = 6.7) while 6/86 retained complete *inlA* (V = 46.7), producing a score range of 16.1 to 32.1 within a CC treated as a homogeneous group by conventional typing. CC9 (n = 38): heterogeneity in persistence markers (*qacH*: 52.6%; *bcrABC*: 23.7%) with P-Scores of 51.2 to 86.4 reflecting qualitatively distinct degrees of environmental adaptation. CC5 (n = 120 global): geographic heterogeneity in *inlA* (100% complete in USA vs 100% truncated in Norway).

### 3.7 Genome-environment discordance: distinguishing intrinsic persistence from environmental opportunity

The Fagerlund dataset enables a validation that the three remaining datasets cannot offer: direct comparison between trophic classification based exclusively on the genome (Level 1 P-Score: genetic markers without temporal metadata) and trophic classification integrating longitudinal surveillance data (Level 1+2 P-Score: genetic markers + temporal monitoring data). The discordance between both classifications constitutes a biological signal with direct interpretive value.

**Table 8.** Trophic migration matrix: genome-based classification (Level 1) vs. surveillance-informed classification (Level 1+2) in the Fagerlund dataset (n = 513)

| Genome-based ↓ | Surveillance → | Amphitrophic | Nosotrophic | Saprotrophic | Unassigned | Total |
| --- | --- | --- | --- | --- | --- | --- |
| Amphitrophic |  | 151 | 0 | 0 | 0 | 151 |
| Nosotrophic |  | 60 | 19 | 0 | 0 | 79 |
| Saprotrophic |  | 0 | 0 | 45 | 0 | 45 |
| Unassigned |  | 123 | 0 | 78 | 37 | 238 |
| Total |  | 334 | 19 | 123 | 37 | 513 |

Three patterns emerge from this matrix:

1. Complete concordance for genome-based amphitrophic and saprotrophic isolates. All 151 isolates classified as amphitrophic by genome analysis remained amphitrophic with temporal data (100% stable). All 45 genome-based saprotrophic isolates remained saprotrophic (100% stable). The genome correctly predicts their trophic strategy: when a strain simultaneously carries virulence and persistence determinants, or when it has performed the irreversible genomic trade-off, longitudinal data confirm what the genome anticipates.
2. Nosotrophic→amphitrophic migration: persistence without genetic tools. 60 isolates classified as nosotrophic by genome analysis (V > 65, P_genetic < 35) migrated to amphitrophic when temporal data were incorporated (P_surveillance ≥ 40). These 60 isolates correspond exclusively to CC1 (n = 30, clone MF7036 in factory S2, 2018–2019) and CC14 (n = 30, clones MF3939 and MF7663 in factories S2, M1, and S3, 2011–2020). All 60 are persistent according to the Fagerlund criterion (detection of the same clone in the same facility across ≥2 distinct calendar years). Their genetic profile is unequivocal: 0% SSI-1, 0% *qacH*, 0% *bcrABC*, P_genetic = 23.4 ± 1.6. They carry no recognized persistence marker. Yet they persist: P_surveillance = 78.6 ± 8.1. The environment is sustaining these hypervirulent clones, not their genetics.
3. Resolution of unassigned isolates. Of the 238 isolates without genome-based trophic assignment, 123 (51.7%) migrated to amphitrophic and 78 (32.8%) to saprotrophic with temporal data, leaving only 37 (15.5%) unassigned. Longitudinal surveillance data resolved the genomic ambiguity for 84.5% of previously indeterminate cases.

Overall: 261/513 isolates (50.9%) changed trophic category upon incorporation of temporal data. Of those that changed, 92.0% were persistent (240/261), compared to 62.7% of the concordant (158/252). The discordance between genome-based and surveillance-informed classification is itself a predictor of persistence (χ^2^ = 61.4; p < 0.001).

## 4. Discussion

### 4.1 Multi-dimensional profiling reveals biological patterns inaccessible through conventional typing

The multi-dimensional scoring approach described here addresses a structural limitation of single-metric characterization through four methodological contributions: quantitative integration of independent biological dimensions into a continuous score; empirically calibrated weights reflecting the relative importance of each dimension; comparative normalization across contexts; and dual calibration that adapts the analysis to different ecological scenarios.

Complementarity with cgMLST is illustrated by a concrete case: within CC9 in the Fagerlund dataset, isolates with AD ≤ 7 (epidemiologically indistinguishable by ECDC criterion) exhibit P-Scores of 51 to 86 depending on the presence/absence of *qacH* and *bcrABC*. For outbreak tracing, both subgroups are correctly identified as the same clone. For understanding their ecology, they represent functionally distinct profiles: the former carries acquired disinfectant resistance, the latter remains susceptible to conventional biocides. This intra-clonal functional granularity is inaccessible from cgMLST — not because it fails, but because it was designed for a different question.

### 4.2 Amphitrophism: a third ecological strategy

The finding of greatest conceptual relevance is the identification of the amphitrophic strategy, which represents 39.1% of the analyzed population and constitutes, to our knowledge, the first quantitative demonstration at the population scale that the virulence-persistence trade-off does not operate as an absolute paradigm in *L. monocytogenes*.

It is essential to contextualize this result. Toledo-Arana et al. (2009) demonstrated functional incompatibility between PrfA-dependent (virulence) and SigB-dependent (saprophytic survival) regulatory programs through transcriptomic analysis. Their contribution established the molecular basis of the trade-off and remains valid. However, what they demonstrated was incompatibility of simultaneous *expression* under specific conditions, not impossibility of simultaneous *carriage* of both genetic modules. Amphitrophic lineages have resolved this apparent contradiction: they carry the determinants of virulence (functional *inlA*) and environmental survival (SSI-1), expressing them conditionally depending on ecological niche — the SigB program in the environment, PrfA in the host.

This contrasts with saprotrophic lineages (CC9, CC121), which have performed the genomic trade-off irreversibly: PMSCs in *inlA* physically eliminate the capacity to produce functional InlA, a loss not reversible transcriptomically.

Notably, Møretrø et al. (2024) independently found that CC7 — the prototypic amphitrophic lineage — exhibits *in vitro* virulence not significantly inferior to the reference strain EGDe, while being simultaneously pervasive in processing plants. The convergence between an independent genomic analysis and our multi-dimensional scoring on the same conclusion reinforces the biological validity of this category.

The biological significance is direct: amphitrophic lineages represent the ecologically most versatile group because they do not exhibit the virulence attenuation that in saprotrophic lineages signals adaptation to the environmental niche. A CC7 classified generically as “intermediate” by CC typing masks a profile of elevated environmental persistence with intact invasive capacity — a combination that multi-dimensional profiling quantifies explicitly.

### 4.3 Persistence-resistance co-selection

The P-R co-selection (ρ = +0.591 globally) suggests that environments with chronic persistence histories could constitute antimicrobial resistance reservoirs with implications extending beyond the immediate ecological context. This association, detectable only through simultaneous quantification of both dimensions, warrants further investigation to establish causal mechanisms.

The dual calibration capability enables the same genomic data to be analyzed with weights reflecting different ecological questions — an analytical feature relevant for integrated surveillance where the same genomes are evaluated across environmental and clinical contexts.

### 4.4 Distinguishing intrinsic persistence from environmental opportunity

The genome-environment discordance analysis presented in section 3.7 reveals an analytical capability that, to our knowledge, has not been previously described: the ability to quantitatively determine whether persistence of a clone in an environment is attributable to the strain (genetic equipment for persisting) or to the niche (environmental conditions permitting persistence of non-adapted strains).

The case of CC14 in the Norwegian facilities S2 and M1 is paradigmatic. Clone MF3939 persisted for 8 years (2011–2019) in S2 and clone MF7663 for 8 years (2012–2020) in M1. Both are CC14 — hypervirulent, with functional *inlA* (V = 70.0), associated with severe listeriosis — yet they completely lack persistence markers: 0% SSI-1, 0% *qacH*, 0% *bcrABC*, P_genetic = 24.3. Based on genome analysis alone, these isolates are classified as nosotrophic (specialized in the pathogenic niche, expected transient environmental presence). With longitudinal data, their P-Score rises to 79–88, revealing a persistence the genome does not explain.

The biological interpretation is that when a clone lacking genetic persistence adaptations persists for years in an environment, the persistence reflects niche characteristics rather than strain adaptation. The environment harbors conditions — spatial refugia, substrate availability, reduced competitive exclusion — that permit survival of even genetically non-adapted organisms.

This contrasts with the opposite scenario: a saprotrophic CC9 carrying *qacH* and *bcrABC* that persists reflects genuine genetic adaptation to the environment — the organism is equipped with specific resistance determinants that confer survival advantage.

The scoring approach formalizes this distinction into three interpretive categories:

| Genome-based trophic | Surveillance-informed persistence | Signal | Biological interpretation |
| --- | --- | --- | --- |
| Nosotrophic | High<br>(P_surveillance ≥ 40) | ENVIRONMENT-DRIVEN | Persistence by niche opportunity, not genetic adaptation |
| Saprotrophic | High<br>(P_surveillance ≥ 40) | GENOME-DRIVEN | Persistence by intrinsic genetic equipment |
| Amphitrophic | High<br>(P_surveillance ≥ 40) | DUAL | Both genetic adaptation and permissive environment |

No current molecular typing system — cgMLST, SNP typing, wgMLST — provides this distinction because they operate exclusively on the genome. The contribution of the present approach lies in quantitatively integrating genomic information with longitudinal surveillance data, enabling discrimination between intrinsic capacity (genomic) and environmental opportunity (ecological).

### 4.5 Limitations and methodological considerations

The present work acknowledges the following considerations. Primary validation employs the Fagerlund dataset temporal metadata as a persistence proxy — the best approximation available given the absence of an established biological gold standard, a limitation inherent to the current state of the field (Ferreira et al., 2014; Palaiodimou et al., 2021) and not specific to this work. The three additional datasets, lacking equivalent temporal metadata, provide cross-validation of biological patterns (trade-off, trophic strategies, correlations) in independent contexts, confirming their geographic reproducibility.

The clinical-focused calibration (V:40%, P:20%, C:30%, R:10%), designed for clinical epidemiological analysis, has not been empirically validated with a dedicated clinical dataset with documented clinical outcomes, constituting a priority future work direction.

### 4.6 Future directions

Priority extensions include: validation of clinical-focused calibration with documented outbreak datasets; extension to other pathogens (*Salmonella* spp., STEC, *Campylobacter*) with pathogen-specific component redefinition; extended virulence profiling (25–30 genes) for fine resolution within amphitrophic CCs; and integration of transcriptomic data to directly test the conditional expression hypothesis underlying the amphitrophic strategy.

## 5. Conclusions

Multi-dimensional genomic profiling provides the first empirically validated quantitative approach for characterizing the ecological strategies of *Listeria monocytogenes* at the population scale. With an AUC of 0.933, perfect specificity, and cross-validation across four independent datasets from five countries, this work demonstrates that weighted integration of virulence, persistence, clonality, and resistance captures biological dimensions inaccessible through conventional clonal typing.

The central contribution is the identification and quantification of three ecological strategies — nosotrophic, saprotrophic, and amphitrophic — for which we propose a formal functional trophic taxonomy. The demonstration that 39.1% of the analyzed population adopts the amphitrophic strategy, simultaneously retaining functional invasive capacity and environmental tolerance determinants without detectable genomic sacrifice, constitutes a finding that challenges the dichotomous paradigm of the virulence-persistence trade-off and reveals a previously unrecognized ecological niche. Amphitrophic lineages — invisible to approaches based exclusively on CC classification — represent the ecologically most versatile group, combining environmental persistence with intact pathogenic potential.

A second contribution is the demonstration that comparison between genome-based and surveillance-informed trophic classification enables discrimination between intrinsic persistence (genome-encoded) and environmentally facilitated persistence: when hypervirulent clones lacking persistence markers persist for years in an environment (as with the 60 CC1/CC14 isolates documented in Fagerlund), the discordance signals environmental opportunity rather than genetic adaptation. This analytical capability, inaccessible to systems based exclusively on genomic data, is possible because the approach quantitatively integrates genomic information with longitudinal surveillance data.

These findings have implications for understanding the evolutionary ecology of *L. monocytogenes* in anthropogenic environments and suggest that the species’ adaptive landscape is more complex than the binary clinical-environmental dichotomy implies.

## Data and code availability

The datasets used are available at NCBI under the BioProjects indicated (Table 1). The scoring pipeline source code (computational pipeline, statistical analysis scripts, and figure generation) is available at https://github.com/jgamboa-biotecno/GIF-Framework. The complete scoring system specification is provided as Supplementary Material (Text S1). Additional inquiries may be directed to the corresponding author.

## Acknowledgments

The author thanks Annette Fagerlund, Matthew Stasiewicz, Yu Wang, and Monika Kurpas for the public deposition of their genomic datasets, which made independent validation possible.

---

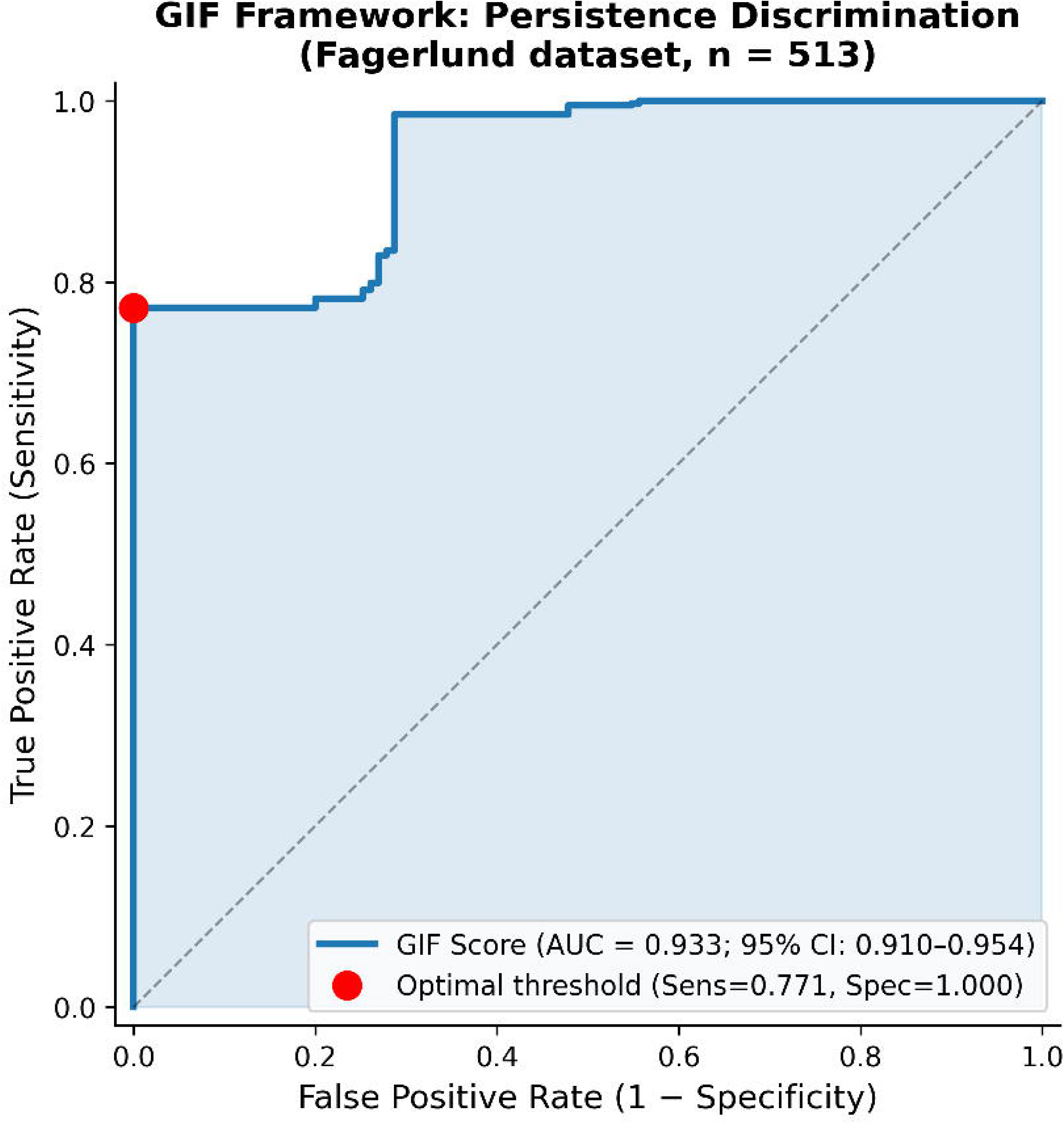

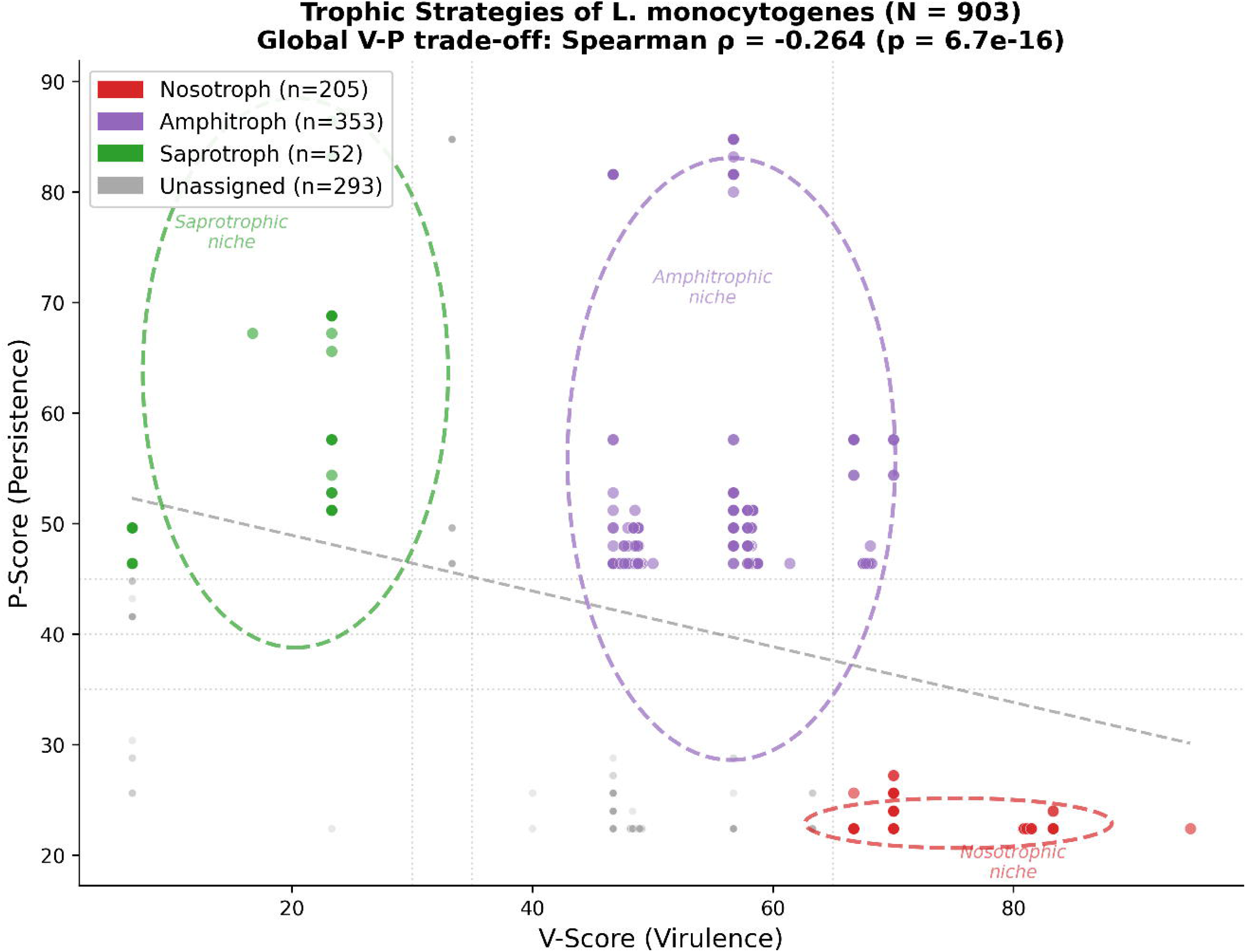

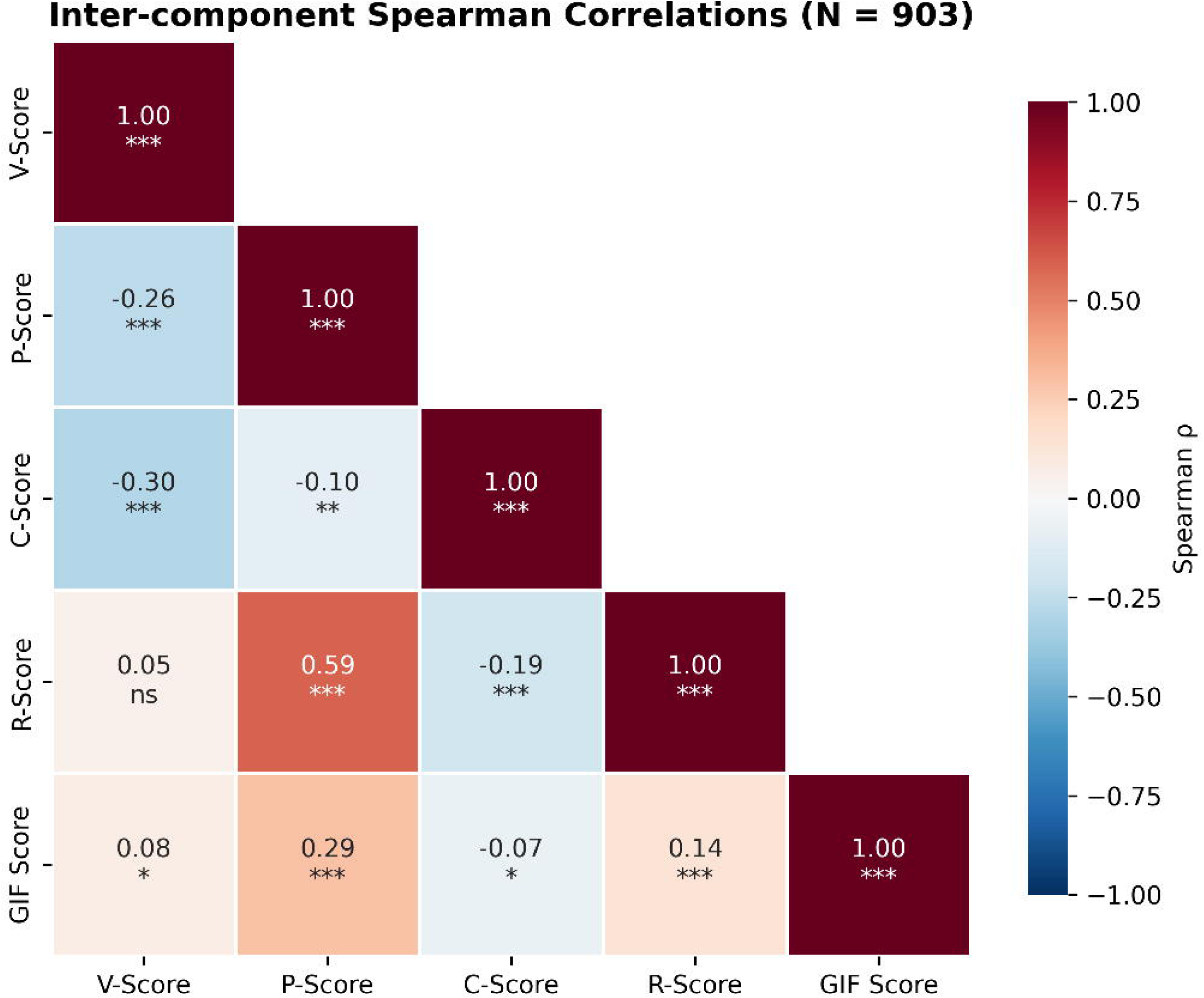

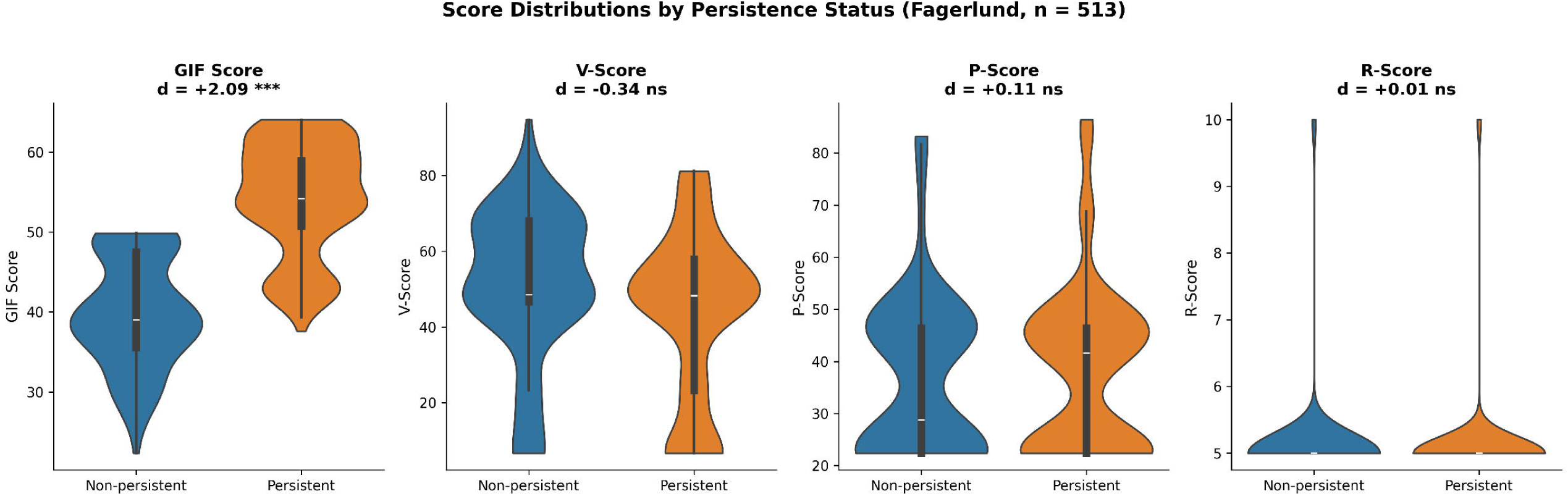

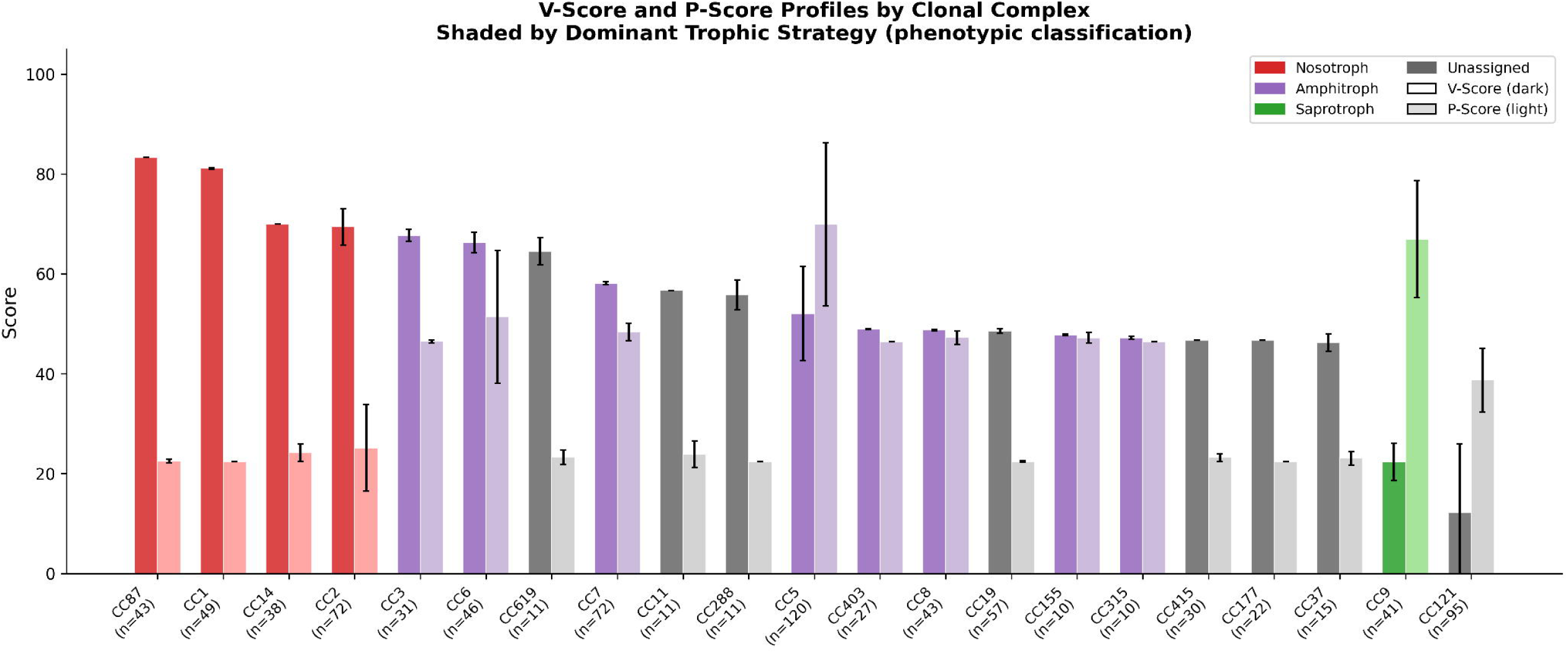

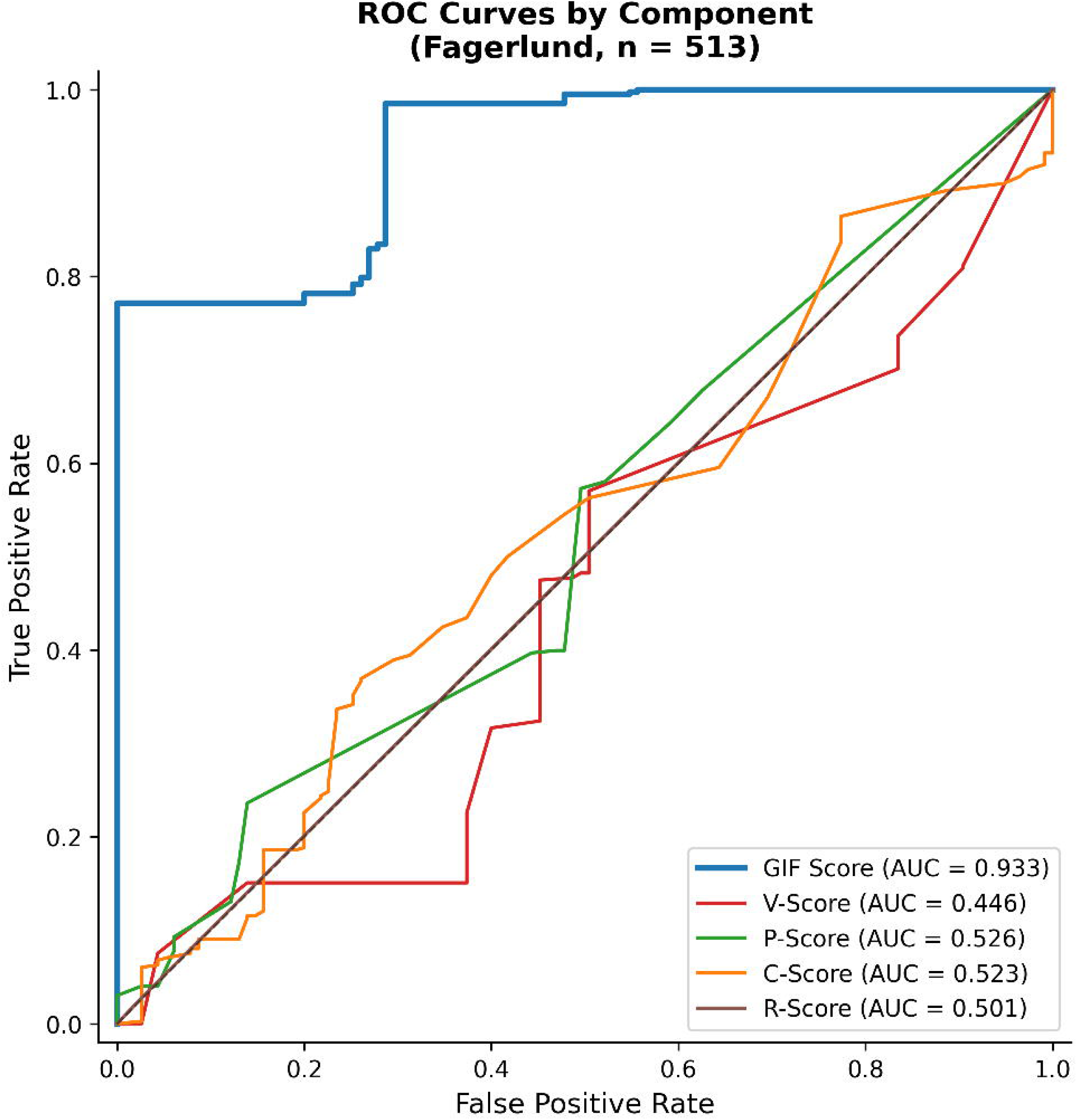

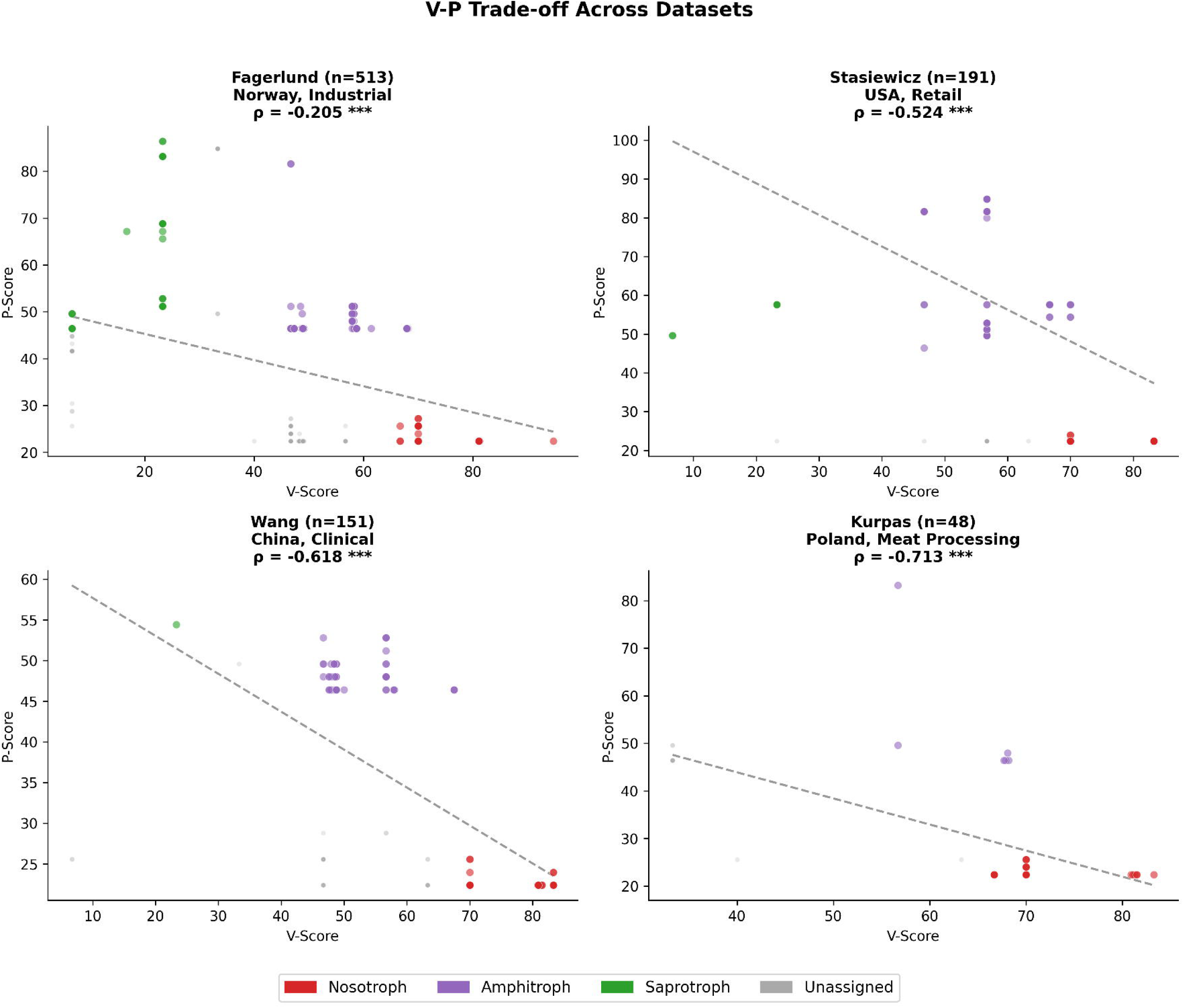

