## Supplementary data for "Amphitrophic *Listeria monocytogenes*: multi-dimensional genomic profiling reveals a third ecological strategy that challenges the virulence-persistence trade-off paradigm"

Javier Gamboa

---

### Text S1. Per-component scoring system — complete detail

#### S1.1 V-Score: Virulence Potential (0–100)

Level 1 — Clonal Complex classification (0–40 points)

| Category | CCs | Points | Clinical frequency | Reference |
|---|---|---|---|---|
| Hypervirulent | CC1, CC2, CC4, CC14, CC87 | 40 | >50% clinical origin | Maury et al. 2016 |
| Intermediate | CC3, CC5, CC6, CC7, CC11 | 20 | 20–40% | Maury et al. 2016 |
| Hypovirulent | CC9, CC121, CC8, CC155, others | 5 | <15% | Maury et al. 2016 |

CC6 classified as intermediate (not hypervirulent) based on updated evidence documenting average virulence for ST6. CC14 and CC87 incorporated as hypervirulent based on documented clinical association.

Level 1 — Genetic proximity to clinical lineages (0–10 points)

| AD_min (cgMLST 1,748 loci) | Points | Interpretation |
|---|---|---|
| ≤7 | +10 | Epidemic cluster (CT) |
| 8–20 | +7 | Related sublineage |
| 21–50 | +4 | Same CC, substantial divergence |
| 51–100 | +2 | Endemic circulation |
| 101–150 | +1 | Sublineage boundary |
| >150 | 0 | Different CCs |

Level 2 — Pathogenicity islands (−10 to +35 points)

| Marker | Points | Condition | Detection |
|---|---|---|---|
| LIPI-1 intact | 0 | Universally conserved | — |
| LIPI-1 deleted | −10 | Deletion in *prfA*, *hly*, or *actA* | BLAST ≥90% id, ≥80% cov |
| LIPI-3 complete | +15 | Lineage I only (valid context) | ≥7/8 genes, ≥90% id, ≥90% cov |
| LIPI-4 complete | +20 | CC4/CC87/CC382/CC619 only | ≥5/6 genes, ≥85% id, ≥80% cov |

Level 3 — *inlA* functional status (−40 to +20 points)

| Status | Criterion | Points | Reference |
|---|---|---|---|
| Complete | ≥780 aa, no PMSCs | +20 | Lecuit et al. 1999 |
| Truncated | 500–779 aa, PMSC at intermediate position | −15 | Nightingale et al. 2005 |
| Severely truncated | <500 aa | −40 | Van Stelten et al. 2010 |
| Not detected | — | 0 | — |

V_raw is normalized to 0–100. Negative values are adjusted to 0.

---

### S1.2 P-Score: Persistence Risk (0–100)

Level 1 — Genetic markers (0–75 points)

*Disinfectant resistance:*

| Marker | Points | Detection | Prevalence Fagerlund (n=513) |
|---|---|---|---|
| *qacEΔ1* | +15 | ≥95% id, ≥90% cov | 0 (0%) |
| *qacH* (Tn6188) | +10 | ≥90% id, ≥85% cov | 76 (14.8%) |
| *bcrABC* (3 genes) | +20 | Each gene ≥90% id, ≥85% cov | 9 (1.8%) |

*Stress tolerance:*

| Marker | Points | Detection | Prevalence Fagerlund |
|---|---|---|---|
| SSI-1 (≥6/9 genes) | +15 | BLAST against lmo0444–lmo0448 | 208 (40.5%) |
| Cadmium tolerance | +5 | cadA/cadC detected | Variable by CC |
| Absence of CRISPR-Cas | +5 | No operon detection | Majority |

Level 2 — Temporal-spatial patterns (0–50 points)

| Metric | Range | Criterion |
|---|---|---|
| Colonization time | 0–30 pts | <4 wk: 0; 4–12: 10; 13–52: 20; >52: 30 |
| Independent detections | 0–10 pts | <3: 0; 3–4: 5; ≥5: 10 |
| Spatial dispersion | 0–10 pts | 1 facility: 0; 2: 5; ≥3: 10 |

Clone definition: ≤7 AD cgMLST (shared SNP reference). Timespan = last − first detection of the clone in the same facility.

Bayesian imputation (without Level 2 metadata): $E[T|G] = (G / G_{max}) \times 25$

Normalization: $P_{raw} / 75 \times 100$, capped at [0, 100].

---

### S1.3 C-Score: Clonality Context (0–100)

Level 1 — Proximity to surveillance database (0–70 points)

| AD_min to surveillance database | Points | Normative reference |
|---|---|---|
| ≤4 | 70 | ECDC multi-country |
| ≤7 | 60 | CT Institut Pasteur; EU 2025/179 |
| ≤10 | 45 | Ruppitsch et al. 2015 |
| ≤20 | 30 | Extended outbreak range |
| ≤50 | 18 | Same sublineage |
| ≤100 | 10 | Endemic circulation |
| ≤150 | 5 | Sublineage boundary |
| >150 | 0 | No clonal relationship |

Level 2 — Clonal expansion (0–30 points)

HC10 cluster size (0–20 pts): ≥100: 20; 50–99: 15; 20–49: 10; 5–19: 5; 2–4: 3; singleton: 0.

Genetic compactness (0–10 pts): AD_median/AD_max ratio within the HC10 cluster. >0.5: 10; 0.3–0.5: 5; <0.3: 0.

---

### S1.4 R-Score: Antimicrobial Resistance (0–100)

Level 1 — Genetic determinants (0–80 points)

| Antimicrobial class | Gene(s) | Points/gene | Maximum |
|---|---|---|---|
| β-lactams | *blaZ*, *blaT*, *blaL* | +5 | 15 |
| Aminoglycosides | *aac(6')-Ie-aph(2'')-Ia* | +10 | 10 |
| Tetracyclines | *tetM*, *tetS*, *tetL*, *tetK* | +10 | 40 |
| Fluoroquinolones (QRDR) | *gyrA*, *parC* mutations | +15 | 60 |

Level 2 — Multi-resistance megaplasmid (0–20 points)

+20 pts if plasmid ≥40 kb carrying ≥3 AMR genes from different categories.

Regional amplification factor: F = max(1.0; P_ref / P_AMR), P_ref = 5%.

---

### S1.5 Contextual calibration

| Parameter | Industrial | Clinical | Justification |
|---|---|---|---|
| w_V | 0.30 | 0.40 | Industrial: chronic exposure > per-event severity |
| w_P | 0.40 | 0.20 | Industrial: persistence is the operational driver |
| w_C | 0.20 | 0.30 | Clinical: outbreak traceability is prioritized |
| w_R | 0.10 | 0.10 | Constant: relevant when present |
| AUC | 0.933 | Pending | — |

---

### Text S2. Trophic classification: quantitative criteria and justification

Trophic strategy assignment is phenotypic — based on V and P scores calculated by GIF for each individual isolate, not on CC membership.

Classification thresholds:

| Strategy | V-Score | P-Score | Threshold justification |
|---|---|---|---|
| Nosotrophic | > 65 | < 35 | V>65 requires hypervirulent CC (40 pts) + complete *inlA* (+20). P<35 indicates absence of persistence markers. |
| Saprotrophic | < 30 | > 45 | V<30 requires penalty for truncated (−15) or severely truncated *inlA* (−40). P>45 indicates presence of SSI-1 and/or qacH/bcrABC. |
| Amphitrophic | ≥ 35 | ≥ 40 | V≥35 requires functional *inlA*. P≥40 requires environmental tolerance markers. Both arsenals retained. |
| Unassigned | Remainder | Remainder | Intermediate profiles without a defined trophic pattern. |

Deterministic consequences of scoring on genomic markers:

The marker profiles observed in each category are a direct consequence of the scoring structure, not a coincidence:

- Every nosotrophic isolate necessarily carries complete *inlA* (verified: 205/205 = 100%).
- Every saprotrophic isolate necessarily exhibits truncated or severely truncated *inlA* (verified: 52/52 = 100%).
- Every amphitrophic isolate carries functional *inlA* (verified: 353/353 = 100%) alongside environmental tolerance markers.
- In the analyzed dataset (N = 903), no isolate exists with complete *inlA* and V < 30, nor with truncated *inlA* and V > 65.

This coherence between quantitative thresholds and underlying biology validates the framework structure.

Implementation:

```
function classify_trophic(v_score, p_score):
  if v_score > 65 AND p_score < 35:
    return "Nosotroph"
  elif v_score < 30 AND p_score > 45:
    return "Saprotroph"
  elif v_score >= 35 AND p_score >= 40:
    return "Amphitroph"
```

```
    else:
        return "Unassigned"
```

---

### Table S1. CC distribution by dataset with dominant trophic strategy

The "Dominant trophic" column indicates the strategy to which the majority of isolates from that CC belong according to individual V-P phenotypic classification. Not all isolates from a CC necessarily share the same strategy.

| CC | Fag | Sta | Wang | Kur | Total | Dominant trophic | Intra-CC heterogeneity |
|---|---|---|---|---|---|---|---|
| CC1 | 34 | — | 10 | 5 | 49 | Nosotrophic | No (100%) |
| CC2 | 7 | 33 | 11 | 21 | 72 | Nosotrophic | Minimal (1% InlA truncated) |
| CC3 | 24 | — | 3 | 4 | 31 | Amphitrophic | No (100%) |
| CC4 | 1 | — | — | — | 1 | Nosotrophic | Insufficient n |
| CC5 | 13 | 83 | 13 | 11 | 120 | Amphitrophic | Yes: 80% amphitrophic, 20% InlA truncated; varies by geography |
| CC6 | 3 | 38 | — | 5 | 46 | Variable | Yes: nosotrophic in Fag/Kur, amphitrophic in Sta |
| CC7 | 69 | — | 3 | — | 72 | Amphitrophic | No (100%) |
| CC8 | 14 | — | 29 | — | 43 | Amphitrophic | No (100%) |
| CC9 | 38 | 2 | 1 | — | 41 | Saprotrophic | No (100% InlA truncated) |
| CC11 | 9 | — | — | — | 9 | Unassigned | Intermediate V, low P |
| CC14 | 34 | — | 4 | — | 38 | Nosotrophic | No (100%) |
| CC19 | 55 | — | — | — | 55 | Unassigned | Intermediate V, low P |
| CC87 | — | 12 | 30 | 1 | 43 | Nosotrophic | No (100%) |
| CC88 | 2 | 4 | — | — | 6 | Amphitrophic | Insufficient n |
| CC121 | 86 | — | 9 | — | 95 | Unassigned | Yes: 86% sev. truncated, 14% InlA complete |
| CC403 | 27 | — | — | — | 27 | Amphitrophic | No (100%) |
| CC415 | 30 | — | — | — | 30 | Unassigned | Intermediate V, low P |
| Others | 67 | 19 | 39 | 2 | 127 | Variable | — |
| Total | 513 | 191 | 151 | 48 | 903 | | |

---

### Table S2. Assembly quality and sequencing methods by dataset

| Parameter | Fagerlund | Stasiewicz | Zhang (Wang) | Kurpas |
|---|---|---|---|---|
| Platform | Illumina MiSeq | Illumina MiSeq | Illumina (via BioNumerics) | Illumina MiSeq (n=40) + NextSeq500 (n=8) |
| Reads | 300 bp PE | — | — | — |
| Assembler | SPAdes v3.10/v3.13 | — | BioNumerics v7.6 | SPAdes v3.11 |
| Contig filtering | <500 bp and cov <5× removed | — | Default BioNumerics | Automatic kmer selection |
| Median coverage | — | 94× (range: 8–360×) | — | — |

| Parameter | Fagerlund | Stasiewicz | Zhang (Wang) | Kurpas |
|---|---|---|---|---|
| Median contigs | — | 26 (range: 12–456) | — | — |
| Median genome size | ~2.9–3.1 Mb (standard *L.m.*) | 3.09 Mb (range: 2.88–3.14 Mb) | ~2.9–3.1 Mb | ~2.9–3.1 Mb |
| %GC | ~38% | ~38% | ~38% | ~38% |
| cgMLST QC | ≥95% of 1,748 loci | ≥95% of 1,748 loci | ≥95% of 1,748 loci | ≥95% of 1,748 loci |
| Annotation | NCBI PGAAP | — | BioNumerics | BIGSdb-Lm |
| QC reference | Fagerlund et al. 2022, Methods | Stasiewicz et al. 2015, Table S3 | Zhang et al. 2021, Methods | Kurpas et al. 2020, Methods |

Note: All genomes were deposited in NCBI and meet minimum quality criteria for cgMLST typing with the Institut Pasteur 1,748-locus scheme. Genome size and %GC values indicated as standard correspond to typical ranges for *L. monocytogenes* reported in the literature (Moura et al., 2016). Stasiewicz et al. (2015) is the only dataset reporting detailed per-isolate assembly quality metrics in its supplementary material (Table S3). Individual per-genome metrics for the remaining datasets can be obtained from the assembly records deposited in NCBI GenBank under the corresponding BioProjects.

---

### Text S3. Strain-facility differential diagnosis: trophic migration methodology

The comparison between genetic trophic classification (based exclusively on Level 1 P-Score: genetic markers) and operational trophic classification (based on Level 1+2 P-Score: genetic markers + temporal monitoring data) was performed exclusively on the Fagerlund dataset (n = 513), which is the only one with real temporal metadata available (factory, isolation year, clonal identifier SNP_ref).

Persistence definition: A clone (identified by its SNP_ref according to Fagerlund et al. 2022) is classified as persistent when detected in the same facility across ≥2 distinct calendar years. This criterion replicates that used by Fagerlund et al. in their original analysis.

Operational P-Score calculation: The Level 2 P-Score incorporates three temporal metrics calculated at the clone level (SNP_ref) per facility (Fag_Factory):

1. *Colonization time* (0–30 points): based on timespan (last − first detection) × 52 weeks/year. <4 weeks: 0; 4–12: 10; 13–52: 20; >52: 30.
2. *Independent detections* (0–10 points): number of clone isolates in the facility. <3: 0; 3–4: 5; ≥5: 10.

3. *Spatial dispersion* (0–10 points): number of facilities where the clone is detected. 1: 0; 2: 5; ≥3: 10.

P_operational = P_genetic_total + P_temporal_real. Normalized to 0–100 with P_norm_max = 75.

Trophic classification: Identical thresholds to the standard classification (Text S2), applied to V-Score (invariant) and P-Score (genetic or operational depending on the scenario).

Migration matrix: Cross-tabulation of the trophic category assigned with genetic P (rows) vs. the category assigned with operational P (columns) for the 513 isolates. Isolates that change category exhibit discrepancy between their intrinsic genetic capacity and their observed temporal behavior.

Operational interpretation of the discrepancy:

| Scenario | Interpretation | Action |
|---|---|---|
| Genetic trophic = nosotrophic, operational trophic = amphitrophic | Strain lacking genetic persistence tools that persists in the plant | The facility has operational deficiencies (niches, hygienic design, L&D protocols) |
| Genetic trophic = saprotrophic, operational trophic = saprotrophic | Concordance: strain equipped to persist, persistence confirmed | Targeted eradication: biocide change, active ingredient rotation |
| Genetic trophic = amphitrophic, operational trophic = amphitrophic | Concordance: strain with virulence + persistence, both confirmed | Maximum priority: dangerous strain in permissive plant |

Association statistic: The association between trophic discrepancy (yes/no) and persistence (yes/no) was evaluated using the $\chi^2$ test with continuity correction.

---

### Table S3. Complete GIF scores for 903 genomes

Available as attached TSV file: GIF_903_genomes_complete_scores.tsv

35 columns for 903 genomes including: identification (Accession, Dataset, MLST_ST, MLST_CC, Lineage), GIF scores (V_Score, P_Score, C_Score, R_Score, GIF_Score), trophic classification (Trophic_Strategy), virulence detail (CC_category, InlA_status, LIPI1/3/4), persistence detail (qacH, bcrABC, SSI1/2), and documented persistence (Fagerlund only).

Fagerlund dataset P-Scores include real temporal data (Level 2) extracted from the supplementary material of Fagerlund et al. (2022). The three remaining datasets use Bayesian imputation.

---

### Supplementary Figure Legends

Figure S1. Per-component ROC curves (Fagerlund, n = 513). Five superimposed curves: composite GIF Score (AUC = 0.933, thick blue line), V-Score (AUC = 0.446, red; inverse correlation confirms trade-off), P-Score (AUC = 0.980, green; near-perfect discrimination), C-Score (AUC = 0.523, orange; neutral), R-Score (AUC = 0.501, brown; neutral). Diagonal line = random classifier.

Figure S2. V-P trade-off across four independent datasets. Four panels: V-Score vs P-Score scatter plots with regression and Spearman ρ. Points colored by phenotypic trophic classification (red: nosotrophic; purple: amphitrophic; green: saprotrophic; gray: unassigned). Significant negative correlation in all four: Fagerlund ($\rho = -0.144$, $p = 0.001$), Stasiewicz ($\rho = -0.524$, $p < 0.001$), Wang ($\rho = -0.618$, $p < 0.001$), Kurpas ($\rho = -0.713$, $p < 0.001$).

---

### Source code

The GIF v1.0 pipeline and analysis scripts from this work are available at:

https://github.com/jgamboa-biotecno/GIF-Framework

The repository includes: - gif_score.py: CLI for GIF-Score calculation from genomic assemblies - analysis/: Statistical analysis and figure generation scripts - README.md: Installation and usage instructions - Dockerfile: Docker image with dependencies (chewBBACA, ABRicate, AMRFinderPlus) - LICENSE: Usage license

---

### Supplementary Material References

Lecuit M, Dramsi S, Gottardi C, et al. (1999). A single amino acid in E-cadherin responsible for host specificity towards the human pathogen *Listeria monocytogenes*. *EMBO Journal* 18(14):3956-3963.

Ryan S, Begley M, Hill C, Gahan CGM. (2010). A five-gene stress survival islet (SSI-1) that contributes to the growth of *Listeria monocytogenes* in suboptimal conditions. *Journal of Applied Microbiology* 109(5):984-995.

Remaining references are found in the main article.
